# Neural correlates of goal-directed and non-goal-directed movements

**DOI:** 10.1101/843011

**Authors:** Naveen Sendhilnathan, Debaleena Basu, Michael E. Goldberg, Jeffrey D. Schall, Aditya Murthy

**Author notes:** Correspondence should be sent to Naveen Sendhilnathan. Raw data and codes will be available upon reasonable request to the corresponding author.

## Abstract

What are the neural correlates that distinguish goal-directed (G) from non-goal-directed movements (nG)? We investigated this question in the monkey frontal eye field, which is implicated in voluntary control of saccades. We found that only for G-saccades, the variability in spike rate across trials decreased, the regularity of spike timings within trials increased, the neural activity increased earlier from baseline and had a concurrent reduction of LFP beta band power.

---

Most movements are goal directed while others, such as fidgets, may not be. However, the neural mechanism that entail these different movements is poorly studied. The macaque frontal eye fields (FEF) in particular has neurons that discharge before visually guided saccades, saccades made in total darkness such as learned saccades or memory-guided saccades, but not before spontaneous saccades in total darkness^1^. Here we discovered that when monkeys make saccades that have no obvious goal in a lit environment, FEF movement and vis-mov neurons do, in fact, discharge. We asked if these seeming non-goal-directed saccades made in the light were actually made to a goal that we did not discern, or if there were differences in neural activities that distinguished between non-goal-directed (nG) and goal directed (G) saccades. We studied two characteristics of neural response not directly visible in the firing rate but which precede movements: a decrease in neural response variability^2^ and a decrease in local field potential beta oscillatory activity^3,4^. Previous studies have shown that decreases in response variability are correlated with attention^5^, preparation of visually guided saccades^2^, the onset of a visual stimulus^6^, etc; and decreases in beta power have been correlated with motor preparation, and inhibitory control^7,8^ among other processes^9^. Nevertheless, despite these efforts, their roles in goal-directed versus non-goal-directed movement generation have not been studied. Here we show that although FEF saccade-related neurons do discharge before nG-saccades, their neural activity was not associated with decreases in either neural variability or beta oscillations, unlike for G-saccades.

We trained two monkeys to perform a visually-guided saccade task where the monkeys made saccades to locations instructed by a prior target presentation. We presented only one target in ~30% of the trials, referred to as ‘no-step trials’ (**Fig 1A**) while in the rest ~70% of trials we presented two sequential targets, referred to as ‘step trials’ (**Fig 1B; Fig S1A** see methods for details). The monkeys either made a single or two sequential visually-guided saccades respectively to earn a liquid reward. The monkeys performed no-step trials with much higher accuracy compared to step trials (**Fig S1C**). We only analyzed correct trials.

**Figure 1:**
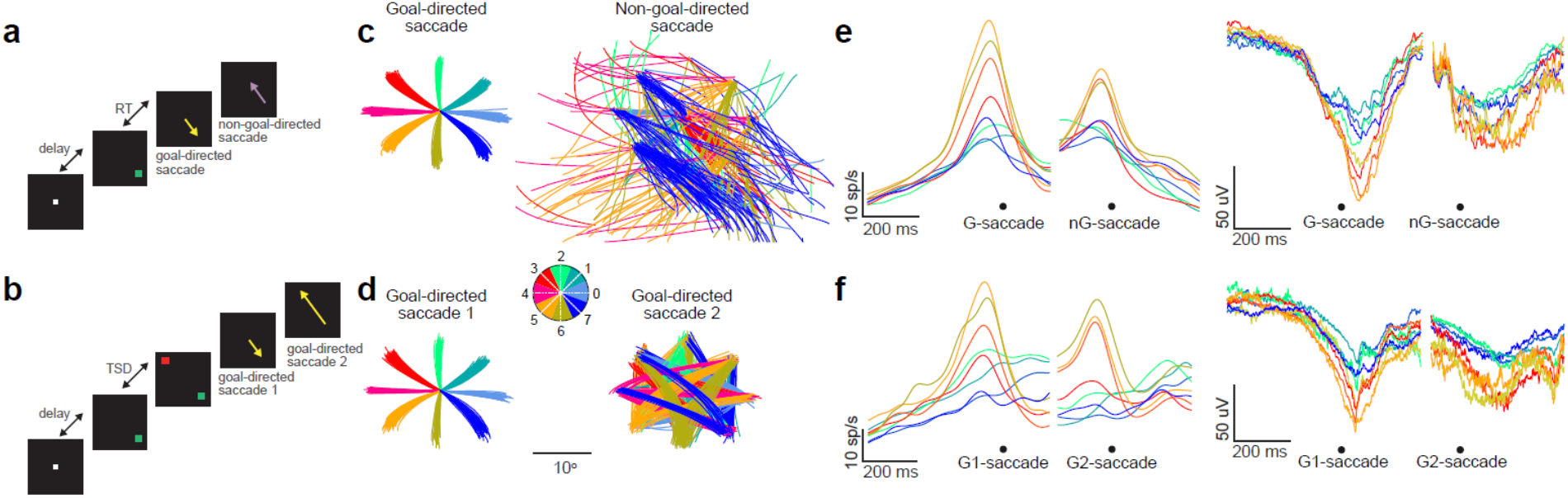
Task, Behavior, and electrophysiology. **a.** Visually guided saccade task - no-step trials: The monkey fixated on a central white square fixation spot on a dark background. Following a variable time delay, a peripheral green target appeared and the monkey made a single G-saccade (yellow arrow) to this target location as soon as possible. The monkey fixated on the target for 200 ms and then was free to move his eyes. He often made task-irrelevant nG-saccade(s), shown in purple, usually back to the fixation spot. **b.** Visually guided saccade task - step trials: The monkey fixated on a central white square fixation spot on a dark background. Following a variable time delay, two peripherals targets (green and red) appeared sequentially and the monkey made two sequential G-saccades (yellow arrow) to these target locations as soon as possible. The monkey fixated on the final target for 200 ms and then was free to move his eyes. **c.** The saccade trajectories from two representative sessions for the G-saccades (left) and nG-saccades (right) in the no-step trials. The colors represent the position ‘to’ which the saccades were irrespective of the starting position (refer to the inset for the colormap legend). **d.** Same as above but for the two sequential G-saccades in step-trials. **e.** Mean population neural activities (left) and mean population LFP activities (right) for no-step trials, for all the positions indicated in C. Same color scheme applies as in C. **f.** Same as F but for two sequential G-saccades in step trials.

In this study, we defined G-saccades as those tasks relevant, visually-guided, reward driven saccades that were instructed by a prior target presentation. We defined nG-saccades as those saccades that were neither visually guided nor instructed by a prior target presentation and thus were task irrelevant and were not rewarded upon execution (these mostly occurred during the inter trial interval period). Thus, monkeys first made either a single G-saccade in the no-step trials or two sequential G-saccades (G1 and G2) in step trials and then made none or several nG-saccades in either trial types, before making a return saccade (R) to the fixation point at the center of the screen to initiate the next trial. These return saccades, were not instructed by a saccade target and were not rewarded immediately; however, they could be considered as G-saccades because they were necessary to initiate the next trial. We only analyzed two consecutive pairs of saccades in every trial: either G immediately followed by nG or return saccades in no-step trials (**Fig 1C**) or two sequential G saccades in step trials (**Fig 1D**) that were matched in inter-saccade-intervals (see methods).

We analyzed simultaneously recorded neural spiking and LFP from 34 visuomotor and 38 movement FEF neurons^3,10^ (**Fig S2**). The vector-averaged spikes and local field potential (LFP) waveforms for nG-saccades were similar to the G-saccades (**Fig 1E, F**). We measured the changes in neural variability across trials using Fano factor (FF): the variance in spike counts across trials divided by the mean across-trial firing rate^2,6^. We controlled for the effect of changes in the mean firing rate on FF by matching the average across-trial firing rate distributions across time bins based on the algorithm developed by Churchland et al^6^. Although the FEF neurons fired with an increased discharge rate during both G and nG saccades (**Fig 2A**), the FF decreased only for the G-saccades (**Fig 2B-C**, P = 1.2997E-07 t-test)^2,11,12^ while it did not significantly change for the nG-saccades (**Fig 2B-C**, P = 0.7435; t-test). Interestingly, since the return saccades could be considered goal-directed in nature, the neural variability decreased for return saccades as well (**Fig S3**).

**Figure 2:**
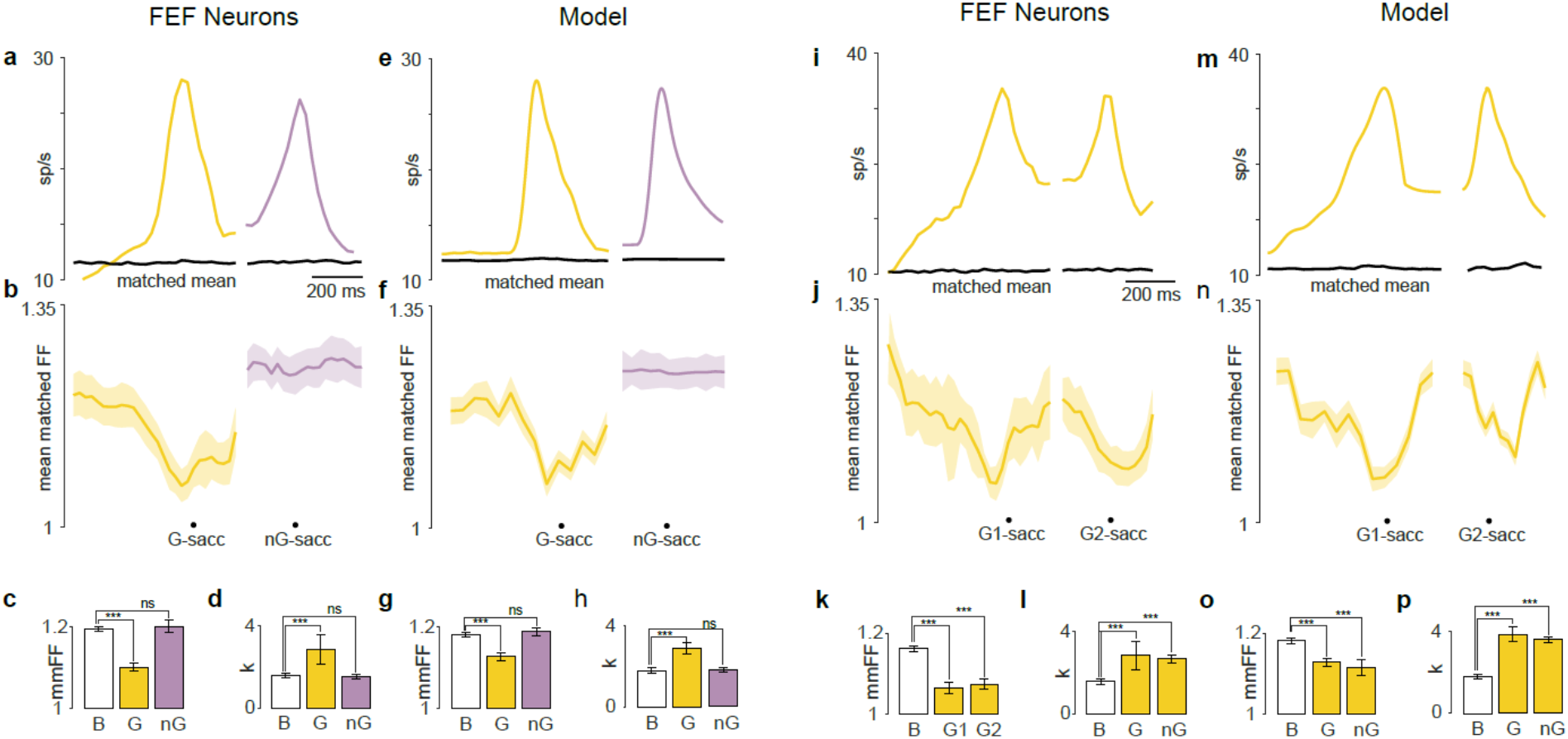
Neural variability decreased only for G-saccades. **a.** Mean neural activity for G-saccades (yellow) and nG-saccades (purple) in no-step trials. The corresponding mean-matched neural activities are shown in black. **b.** Mean matched fano factor for G-saccades (yellow) and nG-saccades (purple). **c.** Quantitation of mean-matched fano factor in B. *** means P = 1.2997e-07, t-test; ns means P = 0.7435, paired t-test. **d.** Within trial variability measured by k was significantly higher for G-saccades (*** means P = 1.8700e-06; ranksum test) than for nG-saccades (ns means P = 0.8972; ranksum test). **e.** Simulated neural activity for G-saccades (yellow) and nG-saccades (purple) from a gamma distribution with k values from D. The corresponding mean-matched neural activities are shown in black. **f.** Mean matched fano factor for the simulated G-saccades (yellow) and nG-saccades (purple) neural activities. **g.** Quantitation of mean-matched fano factor in F. *** means P =2.0127e-04, t-test; ns means P = 0.7324, paired t-test. **h.** Within trial variability measured by k obtained by the simulated spike trains shown in E, was significantly higher for G-saccades (*** means P = 4.3427e-09 ranksum test) than for nG-saccades (ns means P = 0.6342; ranksum test). **i.** Same as A, but for two sequential G-saccades in step trials. **j.** Same as B, but for two sequential G-saccades in step trials. **k.** Same as C, but for two sequential G-saccades in step trials. P =4.400E-03, t-test for G1 saccades and P = 3.9928E-05, t-test for G2 saccades. **l.** Same as D, but for two sequential G-saccades in step trials. P = 1.076E-03; ranksum test for G1 saccades and P = 9.601E-03; ranksum test for G2 saccades. **m.** Simulated neural activity for two sequential G saccades from a gamma distribution with k values from L. The corresponding mean-matched neural activities are shown in black. **n.** Mean matched fano factor for the simulated G-saccades’ neural activities. **o.** Quantitation of mean-matched fano factor in N. P = 3.5645e-05 for G1 saccades and P = 5.4534e-06 for G2 saccades. **p.** Within trial variability measured by k obtained by the simulated spike trains shown in M, was significantly higher for G1 saccades (*** means P = 3.2455e-04 ranksum test) and G2 saccades (ns means P = 4.4565e-04; ranksum test).

The Fano factor describes differences in variability across trials. Additionally, the spike timings within single trials were also less variable during G-saccades than during nG-saccades. To show this, we fit a gamma distribution to the ISI distribution of spike timings during the G-saccades and estimated the shape parameter, ‘k’ that described the variability of spike timings^13^ (see methods). Higher k values represent lower spiking variability within single trials, with k=1 describing a Poisson process. We found that the ISI distribution for G-saccades was characterized by a significantly higher k value than that for nG-saccades (**Fig 2D**, P=2.5712e-06; ranksum test). We then used a previously developed computational model^14^, where we modeled the changes in across trial variability as a consequence of changes in spike time variability within individual trials. Using the experimentally obtained k values, our model predicted a reduction in FF in the G-saccade but not for the nG-saccade, consistent with the experimental observation (**Fig 2E-H**).

Absence of changes in neural variability in the second saccade was not due to sequential planning of saccades (**Fig 2I**). We found decreases in mean-matched FF (**Fig 2J-K**; G1: P=4.400E-03, t-test; G2: 3.9928E-05, t-test) and higher k value (**Fig 2L**; G1: P = 1.076E-03; ranksum test; G2: P = 9.601E-03; ranksum test) for both sequential G-saccades in step trials. Again, our model^14^ captured these data well (**Fig 2M-P**). The neural variability did not decrease for the third (nG) saccade after the two sequential G-saccades in the step trials (**Fig S4**). Taken together, we show that both the within-trial variability in spike timings and across trial variability in neural firing decreased only during G-saccades.

Because the nG-saccades were task irrelevant and were not controlled for, they were heterogeneous in direction, amplitude and velocity. Consequently, such variable kinematic and dynamic factors could confound our observation. However, the FF did not differ for saccades that were matched in amplitude, velocity (**Fig S5**) and direction (**Fig S6**; see methods). Furthermore, the time at which the neural activity increased from baseline before the saccade onset was much earlier for G-saccades (−210 ms) than for nG-saccades (−142 ms; test **Fig S7**; see methods). Interestingly, we also found that the return saccades preceded the nG-saccades in a similar temporal fashion (**Fig S2**).

The LFP activity in the FEF during saccade planning reflects the local neural activity rather than an input to saccade planning^3^ and the frequency component of the LFP provides complementary neural signatures that are not readily observable from just the fluctuations in the raw voltage amplitude. Specifically, the beta band (13-30 Hz) power has been linked to movement preparation and execution in several brain regions^3,15–17^ Beta power is reduced before a voluntary movement, reaches minima around the time of movement execution, followed by a phasic rebound^3,16–18^. We therefore tested whether such activity is modulated in G versus nG-saccades. Average LFP activity decreased both during both types of saccades in no-step trials (**Fig 3A**; G-sacc: P = 2.6410E-21, ranksum test; nG-sacc: P = 2.2795E-14, t-test). However, in no-step trials, the beta band (13-30 Hz) decreased in power only during G-saccades^3^ (**Fig 3B-D**, G-sacc: P = 3.9748E-04, ranksum test; nG-sacc: P = 0.7707, ranksum test). The beta power was directionally invariant for both saccades (**Fig 3E**, nG-saccade: P = 0.2173, ANOVA, G-saccade: P = 0.3264, ANOVA). However, for step trials, for both G1 and G2 saccades, the LFP activity decreased (**Fig 3E**; G1-sacc: P = 2.9855E-09, ranksum test; G2-sacc: P = 4.18813-11, ranksum test), and the beta power was suppressed (**Fig 3F-G**, G1-saccade P = 7.39339E-03, t-test; G2-saccade P = 3.2233E-03, t-test). The beta power was again directionally invariant for both saccades (**Fig 3H**, G1 saccade: P = 0.6234, ANOVA, G2 saccade: P = 0.4573, ANOVA). Taken together, the activity of beta band decreased only for G-saccades and not for nG-saccades.

**Figure 3:**
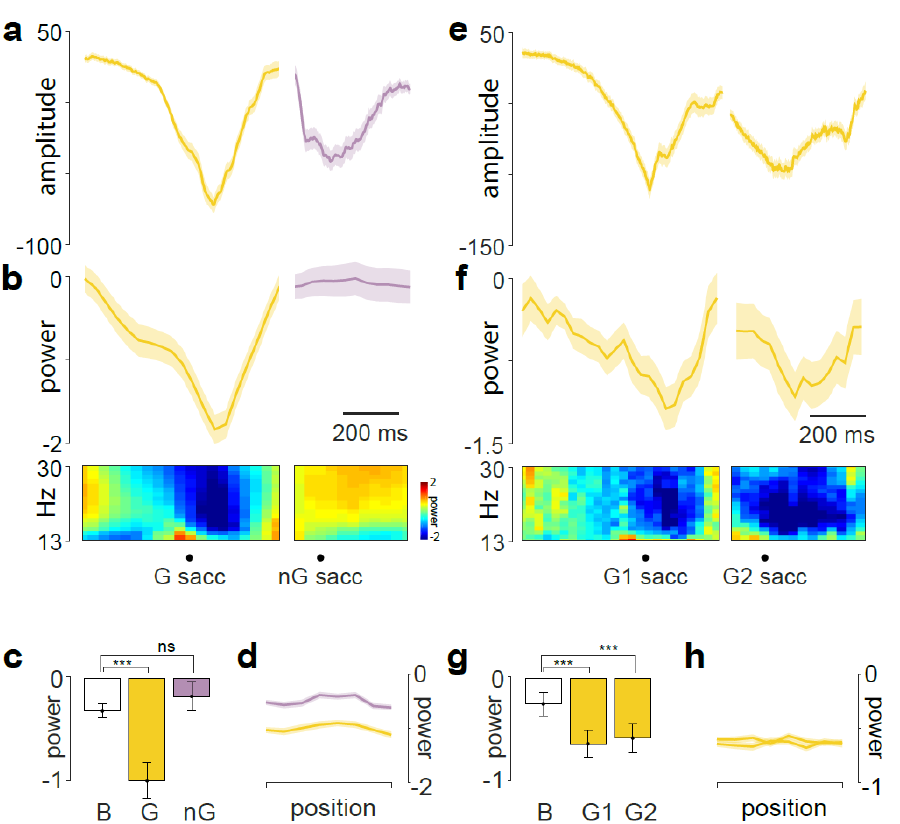
LFP Beta power decreased only for G-saccades. **a.** Mean LFP activity for G-saccades (yellow) and nG-saccades (purple) in no-step trials. **b.** Top: mean Beta band power for G-saccades (yellow) and nG-saccades (purple) in no-step trials. Bottom: Spectrogram showing the beta band modulation. **c.** Quantitation of beta power from B. *** means 3.9748E-04 ranksum test; ns means 0.7707, ranksum test. **d.** Lack of modulation of beta power with positions for G-saccades (yellow) and nG-saccades (purple) in no-step trials. **e.** Same as A but for two consecutive G-saccades in the step trials. **f.** Same as B but for two consecutive G-saccades in the step trials. **g.** Quantitation of beta power from F. B-G1 *** means 7.3939E-03, t-test; B-G2 *** means 3.2233E-03, t-test. **h.** Lack of modulation of beta power with positions for two consecutive G-saccades in the step trials.

In summary, although FEF drives saccades, the neural signatures of G-saccades significantly differed from nG-saccades. Attention can decrease the variability of neural responses, and decrease the shared variance between neurons^19^. In our task, the first saccade was attentionally driven while the second saccade was attentionally driven only on the step trials and for return saccades. The lack of change of neural variability during the non-return second saccades of no-step trials suggests that these saccades were not made modulated by attention. Therefore, our study provides the first conclusive evidence that pre-saccadic neural activity is not necessarily driven by visual attention. Furthermore, a decrease in beta band activity is thought to reflect gradual release of inhibitory control, and is a necessary precondition to execute movements^7^. Our data shows that movement execution can occur in the absence of such a decrease in beta band activity and presumably without a concomitant decrease in inhibition, but selectively for nG-movements. Taken together, our results suggest that only G-saccades are accompanied by cognitive mechanisms such as attention (indexed by decrease in FF) and release of inhibition (indexed by decrease in beta power) and thus add critical constraints to the way we think about saccade generation in the brain.

The question then arises if the FEF activity in nG-saccades is critical for driving them. The FEF movement cells project mono-synaptically to the intermediate layers of the superior colliculus, which are critical for the generation of saccades and presumably drive most saccades. The superior colliculus movement cells fire before all saccades, including spontaneous saccades in total darkness, saccades which are not preceded by FEF activity. Nonetheless it is difficult to postulate that the colliculus is not driven by the FEF for nG-saccades, especially because the FEF activity has a longer presaccadic latency for nG-saccades than the 30 ms minimal latency of the intermediate layers of the superior colliculus^20^. If the FEF induces the superior colliculus to drive nG-saccades in the absence of concomitant decreases in FF and beta, it could be that these non-spike-rate characteristics of neural activity have some function other than the transynaptic transmission of information.

## Acknowledgements

This work was supported by an Intensification of Research in High Priority Areas Grant from the Department of Science and Technology, Government of India; a Department of Biotechnology-Indian Institute of Science (DBT-IISc) partnership programme grant; and institutional support from the Ministry of Human Resource Development. This work was also supported by R21 EY-020631, 1 R01 NS113078-01, and P30 EY-019007 to MEG, PI and the Dana Foundation through the David Mahoney Chair in Brain and Behavior Research, Columbia University. JDS was supported by Robin and Richard Patton through the E. Bronson Ingram Chair in Neuroscience. NS was supported by Kishore Vaigyanik Protsahan Yojana fellowship awarded by the Department of Science and Technology of the Government of India.

## Materials and methods

We performed all our analyses on previously published datasets of frontal eye field neurons^3,10^. Please refer to that study for full details. We briefly describe the experimental procedures and methods here.

### Subjects

Two adult monkeys, J (male, *Macaca radiata*) and G (female, *Macaca mulatta*) were used for the experiments and were cared for in accordance with the Committee for the Purpose of Control and Supervision of Experiments of Animals (CPCSEA), Government of India, and the Institutional Animal Ethics Committee (IAEC) of the Indian Institute of Science.

### Memory-guided saccade (MGS) task

We used this task to classify FEF cells into visual, visuomovement and movement neurons. Details of this task have been described in detail elsewhere^3^. The monkeys were required to fixate on a central fixation point (FP) for a variable amount of time (~300ms) at the start of the trial, following which a gray target appeared briefly at one of the eight equally-spaced peripheral locations on an imaginary circle of radius 12°. The monkeys had to continue fixation on the central FP for a variable delay (1,000 ms ± 15% jitter, sampled from a uniform distribution), following which the FP disappeared (go signal), cueing the monkeys to make a goal-directed memory-guided saccade to the remembered target location, following which the target briefly re-appeared. On successful trials the monkeys received a juice reward. The monkeys were not required to maintain fixation at the target location after the end of the goal-directed saccade. Therefore, they often made none or several non-goal-directed saccades before returning back to the center of the screen in preparation for the next trail.

### FOLLOW task

Details of this task have been described in detail elsewhere^10^. Each trial started with the appearance of a white central fixation point. The task comprised of two types of trials that were randomly interleaved: no-step trials (30%), and step trials (70%). In no-step trials, following fixation for a variable duration (~300 ms), a peripheral green target appeared in one of six equally-spaced peripheral locations (see below) on an imaginary circle of radius 12°. The appearance of the symbol was the go cue for the monkey to make a saccade to the target as soon as possible. In the step trials, after the fixation point appearance, two targets appeared sequentially (initial: green and final: red), with a variable time delay between them, referred to as the target step delay (TSD). We used three TSDs: 16 ms, 83 ms, 150 ms. We flashed the second stimulus almost always in the hemifield diametrically opposite to the hemifield of the first target position. A correct response in the step trials entails making a sequence of two goal-directed saccades: from FP to the first target, and from the first target to the second target. Correct responses were rewarded with liquid reward.

While the MGS task had 8 possible target locations, a restricted set of target positions and steps were used during neurophysiological recording sessions to maximize collection of relevant FOLLOW task data. After response field (RF) identification using the MGS task, typically three target locations centered on the RF were considered to be ‘in-RF’ positions and the three positions diametrically opposite to them were considered to be out of the response field or ‘in-aRF’ positions. The targets in no-step trials and the first targets in step trials could appear in any one of the 6 inside-RF and in-aRF positions. However, the second target in step trials could only appear in one of the three positions diametrically opposite to the position of the first target. Thus, the second target could either step into or out of the response field but never within or adjacent to it. Following successful completion of the task, monkeys made no or several non-goal-directed saccades till they finally made a saccade back to the center of the screen to initiate the next trial. Trials with artifacts including eye blinks, saccades with reaction times less than 100 ms on both tasks were removed prior to further data analysis.

### Data collection

TEMPO/VIDEOSYNC software (Reflective Computing, St. Louis, MO, USA) was used simultaneously with Cerebus data acquisition system (Blackrock Microsystems, Salt Lake City, UT, USA) for data collection. Eye positions were sampled with a monocular infrared pupil tracker (ISCAN, Woburn, MA USA), interfaced with TEMPO software real time. The stimuli were presented on a calibrated Sony Bravia LCD monitor (21 inch; 60 Hz refresh rate) placed 57 cm in front of the monkey. Raw neural signals from 70 neurons were collected using single tungsten microelectrodes (FHC, Bowdoin, ME, USA; impedance: 2 to 4 MΩ) from the FEF on the right hemisphere through a permanently implanted recording chamber (Crist Instrument, Hagerstown, MD, USA). These raw neural signals were acquired at 30000 Hz and was subsampled to 1000 Hz, low pass filtered (from 0 to 150 Hz) to obtain LFP and high pass filtered (300 to 3000 Hz) to obtain spikes.

## Data analysis

### Fano Factor

We calculated Fano Factor (FF) by dividing the variance of spike counts by its mean in continuous bins throughout the trial (with a sliding window of 50 ms shifted by 10 ms). We calculated the mean-matched firing rate using the algorithm developed by Churchland et al.^6^ using a window of length 50 ms, shifted by 25 ms. Each saccade epoch was defined as −200 ms to 100 ms centered on the saccade onset.

### Saccade matching

To match saccades from population *A* to saccades from population *B,* it suffices to find a subpopulation of saccades *A′* such that *A′* ⊆ *B*. Therefore, for each saccade *i* from A (*A_i_*), we took its saccade amplitude and peak velocity and drew a tolerance window around it: ±1° amplitude and ±25°/s velocity. If we found at least three saccades from *B* within every *A_i_*‘s tolerance window, then we classify that *A_i_* as being ‘matched’. By this way, 99.34% of G saccades were matched with G1 saccades but only 50.86% of nG saccades were matched with G2 saccades and the remaining saccades were discarded.

### Matching inter-saccade intervals

The monkeys were required to fixate on the final target, in either trial types, for 200 ms. On average, the monkeys initiated the first nG-saccade ~350 ms after the G saccade in the no-step trials and the monkeys initiated the G2 saccade ~200 ms after the G1 saccade in the step trials (**Fig S1D**). To match the inter-saccade intervals between the two types of trials, we restricted all our analyses henceforth to only pairs of consecutive saccades whose saccade onset times were at least 300 ms apart. The reaction times for the G and G1 saccades were similar individually for both monkeys while the reaction time for G2 saccades was much longer, as expected (**Fig S1E**).

### SPO and SPS calculations

We defined the SPO time as the first time point when the signal significantly differed from the baseline (P < 0.05) continuously for at least the next 20 ms. To calculate SPO, we performed a t-test between the average baseline value and the signal from −400 ms to 200 ms from the start of saccade. This calculation gave a P-value for every millisecond of the data indicating the probability that the signal did not significantly differ from the baseline. Hence, the first time point when the P value fell below 0.05 backwards from the saccade onset and remained below 0.05 continuously for the next 20 ms was taken as the SPO (**Fig S7**).

We defined the SPS time as the first time point when the signals in at least one of the eight positions significantly differed from each other. To calculate SPS, we computed a p-ANOVA between the signals across the eight spatial positions for every millisecond of the data from −400 ms to 200 ms from saccade onset and followed the same steps as mentioned above for SPO (**Fig S7**).

### Gamma distribution fit

The gamma probability density function is a maximum entropy probability distribution and is defined by two parameters: shape (k) and scale (θ). Both these parameters are positive real numbers. The general form of the density function is given by the following equation:

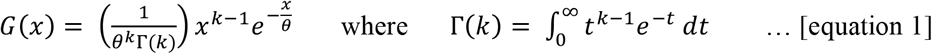

We calculated ISI histograms and fit them to gamma distributions (as shown above) using MATLAB’s gamfit function. This calculates the maximum likelihood estimates of k and θ. After fitting the data, we performed a chi squared goodness of fit test that returned the decision for the null hypothesis that the data comes from a gamma distribution with parameters k and θ, estimated from the data. In specific cases where the data available per analysis was low, this test’s performance declined. Therefore, only in those cases, we visually inspected the fit for confirmation^13^.

### Model simulations

The details of this model are presented elsewhere^14^. Here we describe the model briefly. We simulated 70 neurons with 8 conditions (target positions) per neuron and 20 trials per condition (to match our experimental dataset). For each neuron, four continuous conditions were randomly chosen to have a higher firing rate to the presented target compared to the other four conditions to mimic the RF-aRF property of neurons. We used the gamma probability density function (equation 2) with a shape parameter (*k*, that approximates the inter-spike interval) and scale parameter (θ, that approximates the firing rate) to generate spike trains with a firing rate, *r* given by:

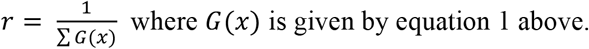

For each simulation, we fixed one of the two parameters: k, θ and we estimated the other parameter iteratively. Finally, we used the mean-matching FF algorithm^6^ to calculate FF on the matched mean.

### Spectrum and Spectrogram

LFP spectra were computed using mtspectrumc and spectrograms were constructed using mtspecgramc functions in Chronux using the multi-taper algorithm^21^. We used five tapers for each analysis and a window length of 300 ms with step size 30 ms to calculate the spectrogram. Spectrograms calculated in this way were normalized by subtracting the log of each value with the log of the baseline power spectrum, in the respective frequency ranges, to get the change in power for each frequency component with respect to time. The frequency range from 13-30 Hz was taken as the beta band for all further analyses.

### Statistics

To check if two independent distributions were significantly different from each other, we first performed a two-sided goodness of fit Lilliefors test, to test for the normality, then used an appropriate t-test; or else a non-parametric Wilcoxon ranksum test. All values in this study, unless stated otherwise, are mean ± s.e.m.

---

**Figure S1:**
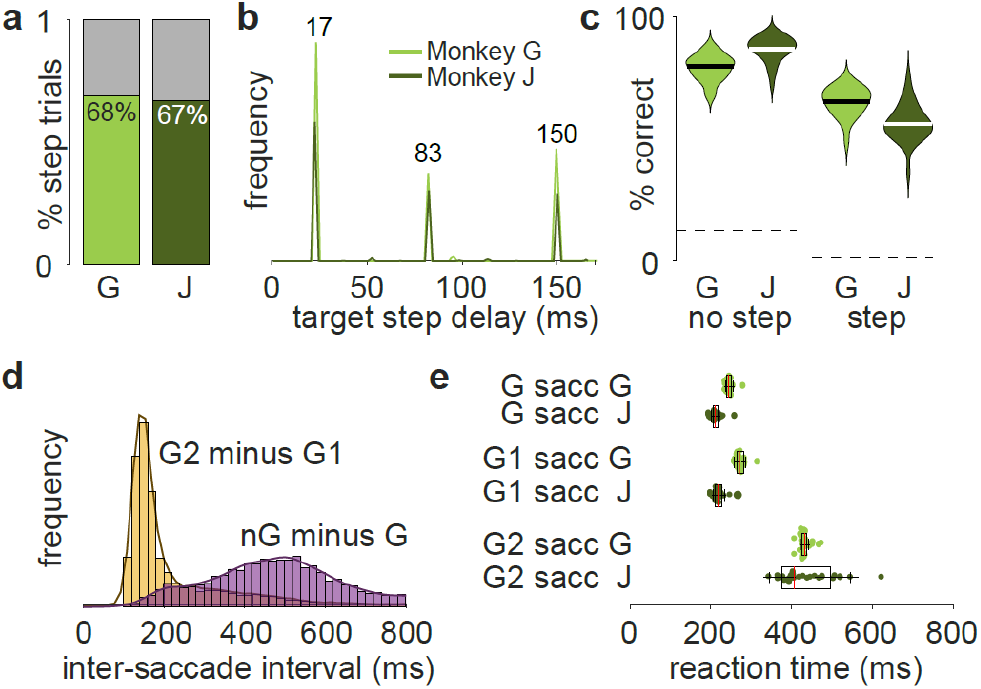
Task and behavior parameters. **a.** Average percentage of step trials (green) for both the monkeys **b.** Target step delay values used for both the monkeys **c.** Percentage of correct trials in the no-step and step trials for both the monkeys **d.** Distribution of intersaccade intervals in the no-step (purple) and step (yellow) trials. **e.** Reaction times for both the monkeys for different saccade conditions.

**Figure S2:**
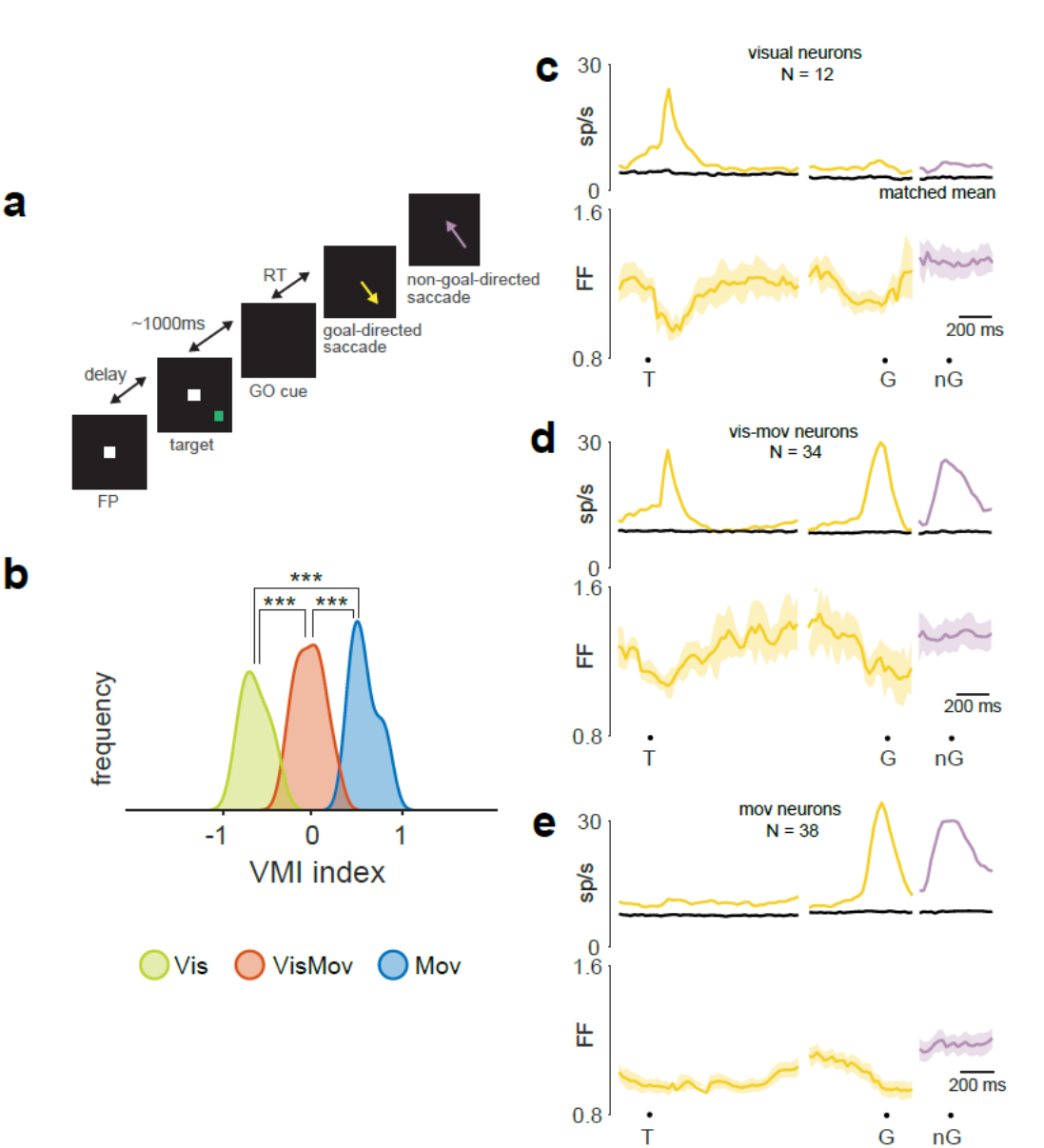
Cell-type dependent changes in neural variability. **a.** Memory guided saccade task: The monkey fixated on a central fixation point on a dark background. Following a variable time delay, a peripheral target appeared briefly on one of eight equally spaced positions on an imaginary circle of eccentricity 12°. The monkey continued to fixate on the central fixation point for 1,000 ms (±15% jitter). When the central fixation spot was extinguished, the monkey made a single saccade (yellow arrow) to the remembered target location. The monkey fixated on the target for 200 ms and then was free to move his eyes. He often made task-irrelevant nG-saccade(s), shown in purple, usually back to the fixation spot. **b.** VMI distributions of the visual (green), vismov (red) and the mov (blue) neurons recorded. **c.** Top: mean neural activity of all the visual neurons (N=12). Bottom: mean matched Fano factor for the visual neurons. **d.** Same as D but for vismov neurons (N =34). **e.** Same as D but for vismov neurons (N =38).

**Figure S3:**
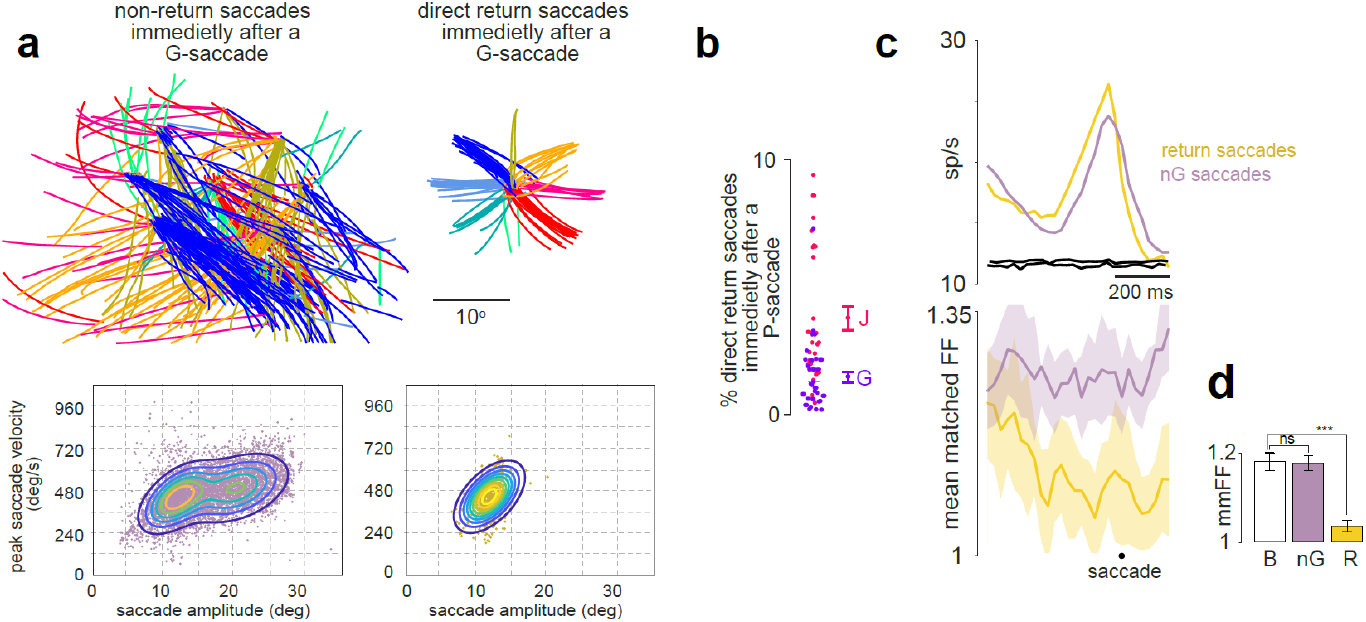
Return saccade and nG-saccade. **a.** Top: Same saccade traces from Fig 1c right but separated based on if it was to an arbitrary position with an arbitrary amplitude (left) or if it was a return saccade. Same color scheme as Fig 1c. Bottom: main sequence for corresponding saccade types across all sessions. Contour map shows the density of data. **b.** Beeswarm plot of percentage of return saccades in each session (each marker is a single session), separated by two monkeys (pink: Monkey J, purple: Monkey G). Average with sem error bars for each monkey is shown to the right. **c.** Top: mean neural activity of return saccades (yellow) and nG-saccades (purple) in no-step trials. black traces are mean-matched firing rates for both conditions. Bottom: mean matched Fano factor for both conditions. **d.** Quantitation of C; *** means P = 8.5872e-06; ns means P = 0.8677;

**Figure S4:**
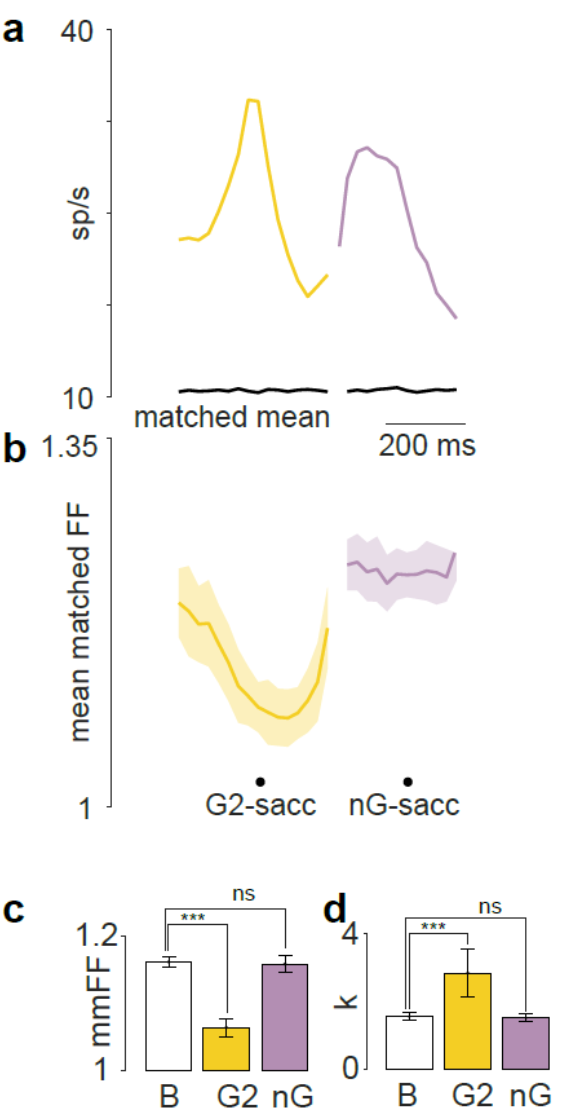
Neural variability decreased for nG-saccade after the two sequential G-saccades in step trials. **a.** Mean neural activity of all the vismov and mov neurons during second G-saccades (yellow) and nG-saccades (purple) in step trials. **b.** Mean matched Fano factor **c.** Quantitation of b; *** means P = 8.929e-05, ttest; ns means P = 0.1316 ttest. **d.** k was significantly higher for G2-saccade (*** means P = 1.233e-04 ttest) than for nG-saccades (ns means P = 0.1443, ransksum test).

**Figure S5:**
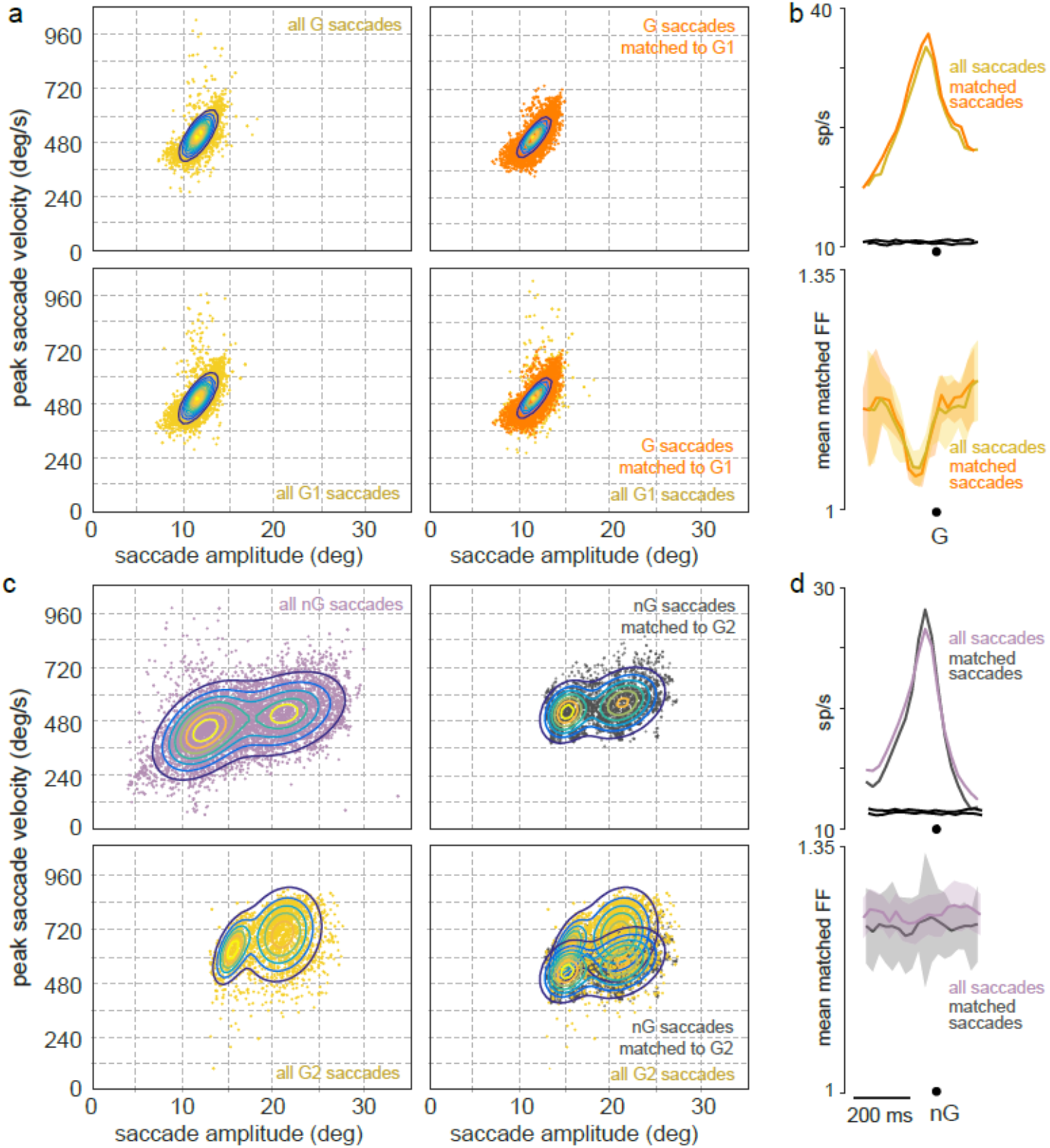
Changes in neural variability are independent of saccade kinematcics. **a.** top left: Main sequence plotted for all the G saccades (yellow markers) in no-step trials. Bottom left: Main sequence plotted for all the G1 saccades (yellow markers) in step trials. Top right: Main sequence plotted for all the G saccades in no-step trials that were matched with G1 saccades in step trials (orange markers). Bottom right: overlay of plots in bottom left and top right. The contour shows the density of data points. **b.** Top: mean neural activity for all G saccades (yellow) and the G saccades that were matched with G1 saccades (orange). Bottom: Fano factor for both these cases. **c.** Same as a, but for nP and G2 saccades. **d.** Same as b, but for nP and G2 saccades.

**Figure S6:**
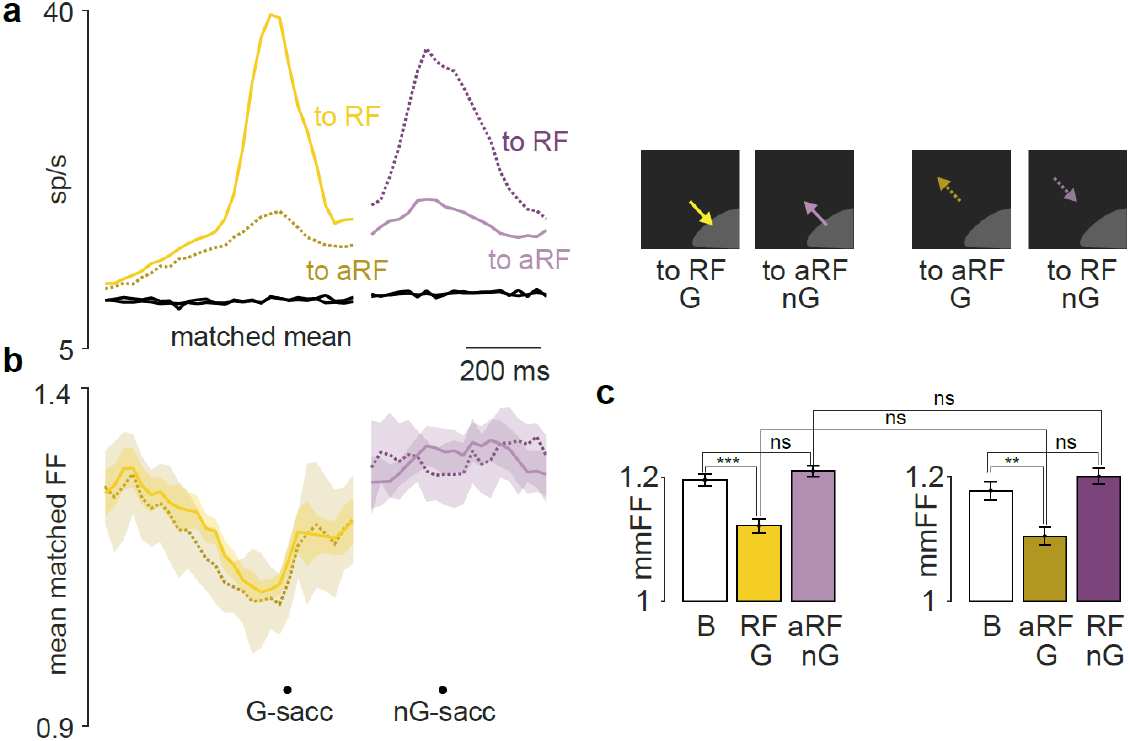
Neural variability was independent of direction of saccade but was dependent on the goal-directedness of the saccade. **a.** Mean neural activity of all the vismov and mov neurons during G-saccades (yellow) and nG-saccades (purple) for saccades into the RF and into aRF (see illustration to the right) in no-step trials. **b.** Mean matched Fano factor for all the vismov and mov neurons during G-saccades (yellow) and nG-saccades (purple) saccades for saccades into the RF and into aRF in no-step trials. **c.** Quantitation of B. G-sacc, B-RF, *** = 7.5486e-05 t-test; nG-sacc B-aRF: ns = 0.3259 ttest; G-sacc: B-aRF, *** = 0.0020 t-test; nG-sacc B-RF: ns = 0.2742 ttest.

**Figure S7:**
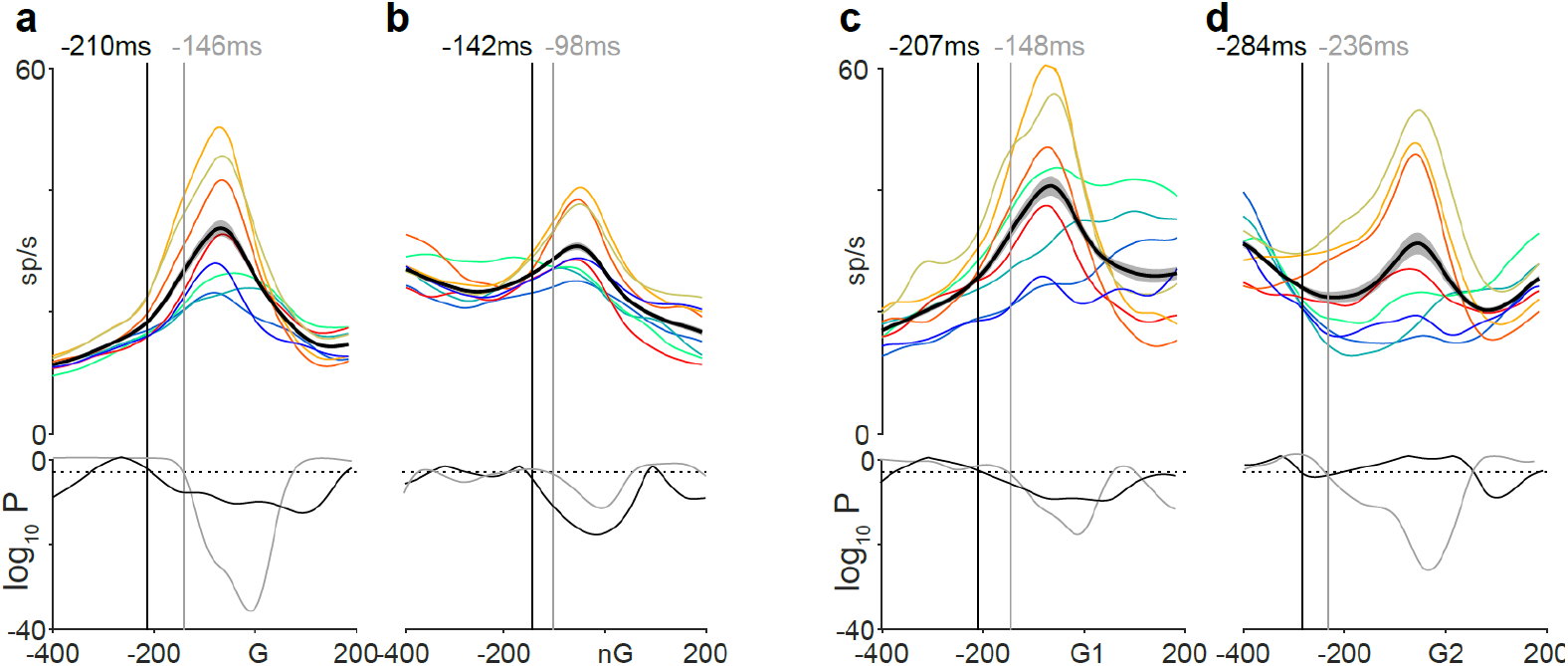
The timing of neural computation for G-saccades was earlier than for nG-saccades. **a.** Top: mean neural activity for each saccade location for G-saccades. Same color scheme as Fig 1. Bottom: The log transformed P value from an ANOVA comparing the mean neural activities across all positions is shown in grey. The time at which this P value started to be significant (fell below the broken line indicating P = 0.05) was taken as the time of SPS (expand) (vertical grey line). Top: The mean neural activity across all positions is shown in black. Bottom: The P value from a t-test comparing the baseline firing with this mean neural activity is shown in black. The time at which this P value started to be significant (fell below the broken line indicating P = 0.05) was taken as the time of SPO (expand) (vertical black line). The SPO was −210 ms and the SPS was −146 ms for G saccades. **b.** Same as a, but for nG saccades in no-step trials. The SPO was −142ms and the SPS was −98 ms. **c.** Same as a, but for G1 saccades in step trials. The SPO was −207 ms and the SPS was −148 ms. **d.** Same as a, but for G2 saccades in step trials. The SPO was −284 ms and the SPS was −236 ms.

## References

1 Bizzi, E. Experimental Brain Research 6, 69–80 (1968).

2 Purcell, B. A., Heitz, R. P., Cohen, J. Y. & Schall, J. D. Journal of neurophysiology 108, 2737–2750, doi:10.1152/jn.00613.2012 (2012).

3 Sendhilnathan, N., Basu, D. & Murthy, A. Proceedings of the National Academy of Sciences of the United States of America 114, 6370–6375, doi:10.1073/pnas.1703809114 (2017).

4 Pesaran, B., Pezaris, J. S., Sahani, M., Mitra, P. P. & Andersen, R. A. J. N. n. 5, 805–811 (2002).

5 Cohen, M. R. & Maunsell, J. H. R. Nat. Neuroscience 12, 1594–1600 (2009).

6 Churchland, M. M., Byron, M. Y. & Cunningham, J. P. Nature Neuroscience (2010).

7 Pogosyan, A., Gaynor, L. D., Eusebio, A. & Brown, P. Current biology: CB 19, 1637–1641, doi:10.1016/j.cub.2009.07.074 (2009).

8 Stancák, A. & Pfurtscheller, G. Cognitive brain research 4, 171–183 (1996).

9 Engel, A. K. & opinion in neurobiology, F.-P. Current opinion in neurobiology (2010).

10 Basu, D. & Murthy. A. bioRxiv, doi:10.1101/390658 (2018).

11 Cohen, J. Y., et al. Journal of Neurophysiology, doi:10.1152/jn.00522.2007 (2007).

12 Chang, M. H., Armstrong, K. M. & Moore, T. The Journal of neuroscience: the official journal of the Society for Neuroscience 32, 2204–2216, doi:10.1523/JNEUROSCI.2967-11.2012 (2012).

13 Maimon, G. & Assad, J. A. Neuron 62, 426–440 (2009).

14 Sendhilnathan, N., Basu, D. & Murthy.A. bioRxiv, doi: 10.1101/635532 (2019).

15 Jasper, H. & Penfield, W. Archivf ür Psychiatrie und Nervenkrankheiten 183, 163–174 (1949).

16 Pesaran, B., Pezaris, J. S., Sahani, M., Mitra, P. P. & Andersen, R. A. Nature Neuroscience 5, 805–811, doi:10.1038/nn890 (2002).

17 Chen, M., Wei, L. & Liu, Y. Scientific reports 4 (2014).

18 Sochůrková, D., et al. Experimental Brain Research (2006).

19 Cohen, M. R. & Maunsell J. H. Nature neuroscience (2009).

20 Wurtz, R. H. & Goldberg, M. E. Journal of Neurophysiology 35, 575–586 (1972).

21 Thomson, D. J. Proceedings of the IEEE (1982).

